# Presurgical language fMRI: Current technical practices in epilepsy surgical planning

**DOI:** 10.1101/279117

**Authors:** Christopher F. A. Benjamin, Isha Dhingra, Alexa X Li, Hal Blumenfeld, Rafeed Alkawadri, Stephan Bickel, Christoph Helmstaedter, Stefano Meletti, Richard A. Bronen, Simon K. Warfield, Jurriaan M. Peters, David Reutens, Monika M. Połczyńska, Lawrence J. Hirsch, Dennis D. Spencer

**Affiliations:** Yale Comprehensive Epilepsy Center, New Haven CT USA; Quinnipiac University School of Medicine, North Haven, CT USA; Northwell Health, Great Neck, NY USA; University of Bonn, Bonn, Germany; University of Modena and Reggio Emilia, Modena MO, Italy; Boston Children’s Hospital, Boston MA USA; The University of Queensland, St. Lucia QLD Australia; University of California Los Angeles, CA USA

**Keywords:** Epilepsy, presurgical, fMRI, language

## Abstract

Little is known about how language functional MRI (fMRI) is executed in clinical practice in spite of its widespread use. Here we comprehensively documented its execution in surgical planning in epilepsy. A questionnaire focusing on cognitive design, imaging acquisition, analysis and interpretation and practical considerations was developed. Individuals responsible for collecting, analyzing, and interpreting clinical language fMRI data at 63 epilepsy surgical programs responded. The central finding was of marked heterogeneity in all aspects of fMRI. Most programs use multiple tasks, with a fifth routinely using 2, 3, 4 or 5 tasks with a modal run duration of five minutes. Variants of over fifteen protocols are in routine use with forms of noun-verb generation, verbal fluency, and semantic decision-making used most often. Nearly all aspects of data acquisition and analysis vary markedly. Neither of the two best-validated protocols were used by more than 10% of respondents. Preprocessing steps are broadly consistent across sites, language-related blood flow is most often identified using general linear modeling (76% of respondents), and statistical thresholding typically varies by patient (79%). The software SPM is most often used. fMRI programs inconsistently include input from experts with all required skills (imaging, cognitive assessment, MR physics, statistical analysis, brain-behavior relationships). These data highlight marked gaps between the evidence supporting fMRI and its clinical application. Teams performing language fMRI may benefit from evaluating practice with reference to the best-validated protocols to date and ensuring individuals trained in all aspects of fMRI are involved to optimize patient care.

## 1.0 Introduction

Neurosurgery is a potentially curative treatment for epilepsy that can be accessed only if the risk that surgery poses to neurological and cognitive function, including language, is known. The validation of language fMRI protocols for predicting language decline after temporal lobe surgery [Bonelli et al., 2012; Sabsevitz et al., 2003], the demonstration of its equivalence or potential superiority to the Wada test (Intracarotid Amobarbital Testing; IAT) with certain protocols [Janecek et al., 2013], and the noninvasive nature of the procedure has led to the widespread adoption of fMRI in neurosurgical planning. In spite of its ubiquitous use, the lack of comprehensive guidelines on precisely how clinical language fMRI should be executed with respect to cognitive design, imaging, procedure, operator training, and interpretation, has led to marked variation in all aspects of the method.

fMRI’s historical development first a research method and the subsequent transition to clinical practice has led to professionals with a wide range of training and skills acquiring, analyzing and interpreting language fMRI data for clinical care. One broad distinction in skill set and expertise is can be made between those working primarily in research and clinical settings. Researchers with doctoral-level specialization in fMRI are typically expert in the method’s strengths and limitations, and, in the context of a focus on null hypothesis significance testing, tend to emphasize avoiding false positive findings. Researchers may have less knowledge of pathology, clinical decision making, and the integration of other test results for clinical care. These factors can result in cautious or tentative application of fMRI when interpreting data pre-surgically for a clinical team. Conversely, clinicians with doctoral-level training in medicine are expert at distilling diverse, complex data into an often binary clinical decision, and are used to making decisions without having near certainty in a given test. They are typically comfortable with taking the available methods and data, and placing the results in a clinical context. They frequently have limited knowledge of the statistical, cognitive, and neuropsychological underpinnings of language fMRI, and its resulting limitations and caveats. In practice, this can manifest as uncertainty when discussing ambiguous or unexpected results (for example, what activation outside Broca’s and Wernicke’s areas represents), the validity of the protocols used, or inconsistent data across paradigms or clinical investigations. An example of this difference between research and clinical approaches is captured in attitudes to the thresholding of clinical fMRI data. While researchers are often hesitant in using non-standardized analysis or unthresholded statistical maps, clinicians may be encouraged by guidelines to vary these [e.g., American College of Radiology, 2014].

Beyond the skill sets that different professions bring to fMRI, the specific decisions made in task design, acquisition, and analysis will also directly impact the results. For example, software packages often implement different solutions in key analysis steps, and fMRI validation studies have used a range of packages such as AFNI [Sabsevitz et al., 2003], SPM [Bonelli et al., 2012] and custom software [Benjamin et al., 2017]. Each step in data preprocessing alters the output; and the literature validating fMRI preprocessing may or may not include explicit realignment of echo-planar imaging (EPI) data, or involvement of explicit smoothing of 2, 10mm or an unspecified amount [Benjamin et al., 2017; Bonelli et al., 2012; Janecek et al., 2013]. The approach to identification of task-related signal also varies, with both the general linear model or a correlation coefficient in use.

Some of the variation in the methods used likely stems from compelling scientific rationale for the use of either of any competing approaches. With respect to the alignment of EPI images, for example, it could be argued that it is optimal to avoid realignment and the attendant further smoothing of data to keep the signal as close as possible to its original (debatably accurate) state. In contrast, it is also reasonable to argue that while this may be feasible in some patients, many others will be unable to remain sufficiently still during scanning, and a reliable method will therefore require realignment. The optimal choices for each variable continue to be subject to debate, though they clearly alter the precise language map obtained [e.g., Benjamin et al., 2018 Figure 3]. While this variation may be less likely to impact lateralization, it will certainly impact localization; and we recently reported that 44% of epilepsy surgical programs use fMRI for this purpose [Benjamin et al., 2018].

The goal of this study was to comprehensively document the current de-facto standards, and the variation therein, for the execution of clinical language fMRI in epilepsy. Such data will allow researchers and clinicians to understand how their methods are applied, how widely adopted their current practices and held beliefs are, and assist in efforts to evaluate and standardize the methods across the clinical fMRI community. We surveyed epilepsy surgical programs who perform language fMRI for clinical care with a detailed questionnaire addressing cognitive design, image acquisition, analysis and interpretation, and practical considerations including personnel and the time involved in completing analysis. Based on the available literature and our own experience we hypothesized marked variation in each of the above across programs.

## 2.0 Methods

This study was reviewed and approved by the Yale Medical Center Institutional Review Board. All respondents provided informed consent.

### 2.1 Survey

A survey centered on fMRI tasks’ cognitive design, image acquisition; data analysis and interpretation; practical issues; and reported accuracy and outcomes was designed (Supplement 1). Questions were generated, reviewed and audited by collaborators in neuropsychology, radiology, neurology and neurosurgery, and reviewed and edited by research consultants (Yale Center for Analytic Sciences). Questions were presented hierarchically on a web-based platform (Qualtrics) with a common set supplemented by follow-up questions as required (e.g., respondents who reported they smoothed data during analysis were then asked what kernel size was used). The survey was designed to be comprehensive while allowing respondents with limited time to skip questions if needed. In these cases respondents were typically required to dismiss a warning prompt to continue. Questions could be returned to and revised throughout.

All respondents assented to the statement “I personally collect, analyze, interpret clinical fMRI data. I use software like SPM (e.g., radiologist, neuropsychologist, imaging scientist etc.).” A second survey, focused on clinicians’ interpretation of fMRI and patient outcomes, was also forwarded to the “epilepsy surgical program [director], or a senior clinician involved in determining patients’ surgical eligibility” with this survey. While these surveys were intended to be paired, most sites did not submit paired responses, and thus the clinical survey is reported separately [Benjamin et al., 2018]. Nine respondents from that manuscript completed both surveys; their responses on accuracy and outcomes are also included here (Section 3.6).

### 2.2 Data collection

Data were collected from 07.17.2015 through 01.15.2016. In the USA we emailed all level 3 and 4 epilepsy centers of the National Association of Epilepsy Centers (NAEC), followed up by telephone in 07.2015, then via email and the American Epilepsy Society (AES) listserv in 11.2015. The NAEC is the major body accrediting epilepsy centers within the USA; levels 3 and 4 centers are surgical programs that complete Wada and/or fMRI. For programs outside the US we adopted a modified snowball sampling approach [Goodman, 1961] to maximize reach and recruitment. We contacted heads of epilepsy organizations, emailed International League Against Epilepsy member boards, and contacted prominent researchers to inform them about the survey. We asked that they identify and forwarded the invitation to epilepsy centers in their regions.

### 2.3 Data analysis

Data were cleaned, with responses entered in error removed (e.g., rare responses that were logically inconsistent). The number of responses per item also varied due to the hierarchical structure of the survey and participants skipping items (which was rare), as noted above. The number of responses per question is indicated throughout the results in square brackets; e.g., [n=X]. Descriptive statistics are presented, and where relevant comparison are made using Fisher’s Exact tests or t tests.

### 2.4 Sample Characteristics

Respondents included sixty-three “analysts” who agreed they “personally collect, analyze, interpret clinical fMRI data [and] use software like SPM (e.g., radiologist, neuropsychologist, imaging scientist etc.).” Of these, 14% reported they also select patients for surgery. When asked further, most respondents reported not being involved in surgical decision making at all (65%) while 35% reported some degree of involvement (8% were Surgical Program director) [n=48 respondents]. Respondents identified as radiologists (29%), neuropsychologists (25%), neurologists (21%), physicists/engineers (10%), neuroscientists (8%), neurosurgeons (2%), MR technologists (2%), and a M.S. in Biomaterials Science (2%) [n=48]. They predominantly worked as both clinicians and researchers (54%), and less often purely as clinicians (29%) or researchers (17%) [n=48].

Responding sites were busy, evaluated children and adults, and completed fMRI in most surgical candidates. Respondents reported evaluating 107 patients annually for surgery (range 10–300; SD 72.6) and 43 receiving surgery (0–151; SD 35) [n=50]. Adults were evaluated at 78% of programs, children at 58% (specifically: 42% evaluated predominantly adults; 36% predominant adults and children; 22% predominantly children). The analysts estimated 65% of surgical candidates at their sites received fMRI for investigating language organization (10–100; SD 28) [n=49], 25% Wada testing [n=39]; 33% extra-operative [n=40] and 27% intra-operative mapping [n=37], and 91% neuropsychological assessment [n=46]. Other methods including magnetoencephalography (MEG), transcortical magnetic stimulation (TMS), gamma activation, cortico-cortico evoked potentials (CCEPS), DTI of the arcuate and other structures, and visual field mapping were used less frequently.

Geographically, analysts were primarily from the US (44%), Australia (11%), Germany (11%), Canada (8%), Italy (6%), France and Switzerland (each 3%), with single respondents from each of Belgium, England, Israel, Scotland, South Africa, Sweden, The Netherlands, and Turkey. Most respondents’ epilepsy programs were affiliated with a university (82%) [n=60].

## 3.0 Results

### 3.1 Cognitive design

A majority of individuals reported use of two or more paradigms (95%), with approximately equal numbers routinely using two (21%), three (20%), four (20%) five (21%), or six or more (14%) language tasks [total n=56 respondents]. Variants of a range of standard neuropsychological paradigms were reported (Table I). The most frequently used was “noun-prompted verb generation” (66%), where respondents generate a verb in response to a (typically visually presented) noun. Verbal fluency (59%), where patients generate as many words as possible in response to a presented letter, was also often reported (“letter-prompted word generation”), as was semantic decision-making paradigm (36%) where, for example, a patient may be presented with two words (e.g., cat-dog, or cat-apple) and decide if they are from the same semantic category.

**Table I:**
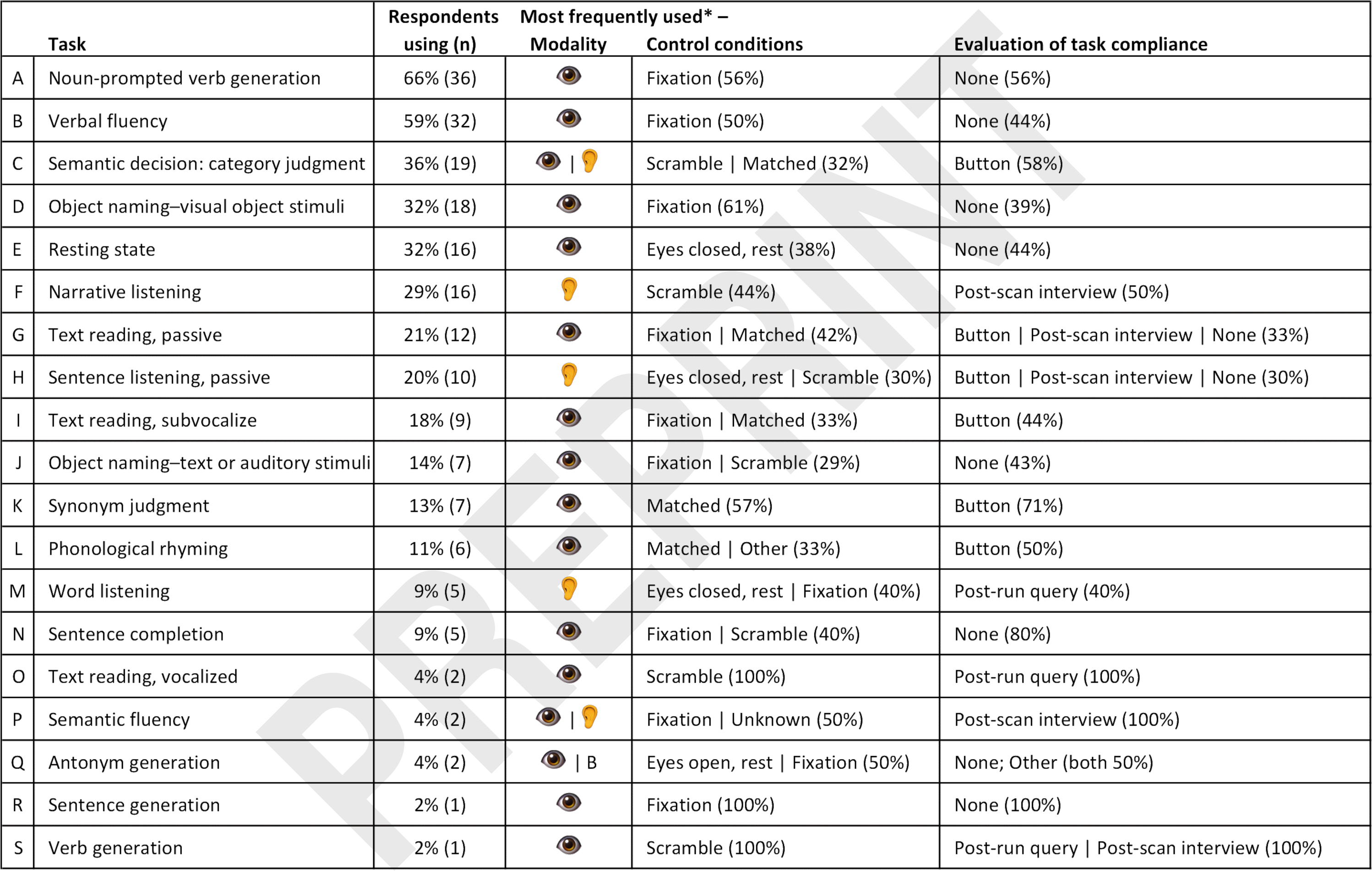
Clinical fMRI paradigm use. <u>(A) Noun-prompted verb generation</u>. A noun is presented aurally or visually, patient asked to think of verbs associated with presented noun, either silently or vocally. <u>(B)</u> <u>Verbal fluency</u> (“Letter-prompted word generation”). A letter is presented aurally or visually, patient asked to think of words that start with the presented letter, either silently or vocally. <u>(C) Semantic decision: category judgment</u>. Two words are presented aurally or visually, patients asked to judge whether words belong to same higher category (e.g. “cat-dog” are in the same category, “cat-apple” are not). <u>(D) Object naming: visual object stimuli</u>. Image of object presented, patient asked to imagine vocalizing name of object silently. <u>(E) Resting state</u>. Patient directed to rest, no response is required. <u>(F) Narrative listening</u>. Auditory stimuli presented, no response is required. <u>(G) Text</u> <u>reading, passive (“Visual language comprehension”)</u>. Text visually presented, no response is required. (H) Sentence listening, passive. Auditory stimuli presented, no response is required. <u>(I) Text reading, subvocalize</u>. Text visually presented, patient asked to covertly imagine vocalizing text silently. <u>(J) Object naming</u>: text or auditory stimuli (“Verbal responsive naming / Description-cued object naming”). Description of object is presented aurally or visually, patients asked to name object. <u>(K) Synonym judgment</u>. Two words are presented visually or aurally, patients asked to judge whether words have similar meanings. <u>(L) Phonological rhyming</u>. Two words are presented visually or aurally, patients are asked to judge whether words rhyme. <u>(M) Word listening.</u> Auditory stimuli are presented, no response is required. <u>(N) Sentence completion</u>. A sentence is presented, the patient generates the final word (multiple respondents noted this was from the Invivo system). <u>(O) Text reading, vocalized</u>. Vocalized text reading text presented visually, patient asked to read text aloud. <u>(P) Semantic fluency</u> (“Category-prompted word generation”). The patient is given a category and names things belonging to that category. <u>(Q) Antonym generation</u>. The patient generates antonyms of presented words. <u>(R) Sentence generation</u>. The patient reads a visually displayed word and makes a sentence that includes the word. <u>(S) Verb generation</u>. Detail unclear; may reflect noun-verb generation. *Most frequently reported response(s) noted. Modality: stimulus modality; B = both. Respondents viewed the task title and elicited a full description by clicking on the title. For further detail, please see Supplement B.

When reviewing the tasks in use, eyes-open rest with crosshair fixation was the most frequent control condition (38%) specified by respondents [n=191 tasks; 54 respondents]. An active control was also frequently reported (17%), as was a scrambled version of the task (15%). Eyes closed rest (14%), eyes open rest without a visual stimulus (9%), and white noise (6%) were also noted. In 10 instances respondents did not know the control condition used; in a further 20, respondents could not categorize the condition. Note that “active” controls could match the task condition in variable ways; for instance, finger-thumb opposition was used as a control for silent visual object naming, verbal fluency, noun-verb generation, and verbal responsive naming in some instances. Stimuli are most often visual (63%), with paradigms less often using auditory (29%) or both visual and auditory stimuli (8%) [n=206 tasks; 54 respondents]. A blocked design is overwhelmingly used (95%) [n=195 tasks; 54 respondents].

Some of the best-studied tasks were identified in a recent, exhaustive review of the evidence supporting the use of language fMRI for the American Academy of Neurology Practice Parameters [Szaflarski et al., 2017]. An approach approximating the approach validated by Binder and colleagues [Sabsevitz et al., 2003], which uses a semantic decision task paired with an active control and uses a laterality index in analysis (whole-brain or regional), was reported by up to 7% of programs. A similar approach to that used by Bonelli [2012], lateralizing language by completing both verbal fluency and noun-verb generation, using a crosshair control and region of interest analysis, was reported by up to 5% of programs.

*Languages imaged.* Sites reported completing language fMRI in a wide range of languages, including English (80%); Spanish (44%); French and German (18%); Arabic, Italian and Turkish (12%); Persian/Farsi (4/6%) and Russian (10%); Hindi (8%); Mandarin (6%); Dutch, Hebrew, Punjabi and Urdu (4%); and finally Bengali, Cantonese, Croatian, Greek, Haitian Creole, Portuguese (European), Slovenian, Somali, Swedish, Swiss German, and Afrikaans (2%) [n=50]. Others also noted use of “Eastern European” languages and, in one case, eight Indian dialects. Among US respondents, most sites imaged in more than one language (68%), with the second most common language at all such sites being Spanish. Other languages imaged in the US included Arabic, Farsi/Persian, French, German, Haitian Creole, Hebrew, Hindi, Mandarin, Punjabi, Somali, Urdu, and a range of other unlisted languages.

### 3.2 Acquisition

*Imaging* was typically completed at 3T (90%) [total n=50 respondents] with isotropic voxels (72%) of 3mm^3^ (41%) [n=31]. Voxel size varied markedly, however, with 1.5mm^3^ [3%]; 2mm^3^ [13%], 3.4mm^3^ [3%], and 4mm^3^ [13%] all in use, and 28% of sites using non-isotropic voxels. All responding sites kept voxel size constant across language EPI runs.

Modal run duration for any given task was 5 minutes 0s (average 4m57s, SD 94s 2m48s-10m) [n=35 respondents/143 tasks]. Nearly all sites used a fixed repetition time (TR) across all runs (94%) [n=34]. Of those using a fixed TR, the modal duration was 3s (47%), with 2s (25%) and 2.5s (22%) also being common [n=32]. The number of runs acquired for any routinely administered language task was typically one (54%), two (11%), or 1–2 runs based on the task or other requirements (14%) [n=35]. Between 3 and 8 runs of tasks were given by the remaining 21% of respondents.

*Patient instruction.* Programs typically had patients practice the tasks prior to scanning (92%), and a majority of sites provided instructions via microphone in scanner (63%) [n=49]. Instructions may be presented on screen in the scanner (45%), and 14% of sites reported incorporating mock scanner practice if needed. Two sites also noted providing patients with written information beforehand, in one case via mail. Another noted that pre-scan practice involved the tasks in their entirety.

*Task compliance* was most often not evaluated (32% of paradigms) [n=261 protocols; 54 respondents]. When analysts evaluated compliance as part of a paradigm, this was most often achieved through a post-scan interview with the patient (24%). Respondents also reported verbally querying patients immediately after the paradigm is run (18%), or using button-press responses in-task (16%). Other methods were used in 10% of cases. For a further 12 protocols it was not known how compliance was evaluated.

### 3.3 Analysis

*Preprocessing.* Programs reported applying standard preprocessing steps with variable frequency.

- *Realignment* within the T2* sequence was completed by 84% of respondents [of a total of n=44];
- *Slice-timing* correction was typically applied (57%) [n=37];
- *Normalization.* Functional data was usually normalized to the patient’s structural image (81%) [n=42], and nearly always to a T1 image such as an MPRAGE (94%). Less frequently a T2 was used as a reference (6%) [n=34]. One program explicitly reported normalizing to a T1 and T2; another noted at times referencing use of a 3D FLAIR instead of an MPRAGE, and a third noted using a T1 gadolinium or T2FSE as needed. Images were not typically normalized to a standard space (e.g., MNI; 18%) [n=39].
- *Smoothing* was typically completed (81%) [n=43], most often with an 8mm kernel (38%) [n=24]. The degree of smoothing varied markedly across sites (2mm [4%]; 3mm [17%]; 4mm [13%]; 5mm [4%]; 6mm [21%] and 10mm [4%]).
- *Motion correction* was typically addressed during analysis (i.e., not on-line; 91%) [n=47], most often statistically (modeling; 47%) while 34% of respondents remove contaminated volumes, in at least some cases, prior to analysis [n=38]. Other reported strategies included rejecting the data and repeating acquisition, and relying on realignment.
- *Other preprocessing* reported included temporal blurring (+/−1/2 TR) and complete “censoring of outlier signal.” A third of respondents did not know if one or more steps were completed [n=48].

*Modeling* was reportedly completed using General Linear Modeling (76%) or a correlation coefficient (26%) [n=42; one site variably used both]. Thresholding was varied on a patient-by-patient basis (“dynamic” thresholding; 79%) [n=47]. Few sites reported using a fixed threshold (19%) or unthresholded maps (2%). The threshold was typically uncorrected (59%) [n=44]. When correction was applied, Bonferroni correction (61%) or the False Discovery Rate (22%) were typically used (other, 17%) [n=18], and applied either voxel-wise (55%) or cluster-wise (45%) [n=11]. One site noted rank-ordering voxel correlation coefficients and thresholding the top 2% brain-wise.

*Software.* Respondents most often used the SPM package for clinical fMRI analysis (27%), with nearly all these sites using SPM8 (75%; SPM5, 8%; SPM12, 17%) [n=48]. Other analysis software reported included InVivo DynaSuite Neuro (Philips) (23%); BrainVoyager (10%); AFNI (Analysis of Functional NeuroImages) (8%); MRIx (4%); Bold MRI Package (Siemens) (4%); Syngo.MR Neuro fMRI (Siemens) (4%); FSL (FMRIB Software Library) (4%); nordicBrainEx (NordicNeuroLab) (2%); Prism (2%); BrainWave (GE) (2%); and other software (8%) such as custom scripts and in one case iViewBOLD (Philips).

*Language areas mapped.* Respondents reported that using fMRI, they routinely systematically map the boundaries of Broca’s area (70%), Wernicke’s area (70%) and basal temporal naming areas (23%), with 28% not routinely and systematically mapping any regions. Other language areas are often identified (32%), with various respondents noting they also map premotor cortex; middle frontal gyrus (“MFG”); anterior insula; supplementary motor area (“SMA”); Exner’s Area; language-related primary auditory and visual cortex; cerebellar language areas; dorsolateral prefrontal cortex (“DLPF”); angular gyrus reading area; visual word form area (reading); and frontal and temporal cortex associated with a second language.

### 3.4 Interpretation

*Interpreting data from different protocols.* In interpreting data from clinical language fMRI protocols, most programs reported considering the maps from different tasks separately and reviewing them visually (73%) [of n=45 respondents]. One third of sites create a single map for interpretation by combining the separate tasks’ maps (e.g., conjunction analysis; 29%). A further 7% create a single map by combining the raw data during analysis. A single one of these approaches is typically used (89%); for instance, overall most programs (62%) interpret data solely by visually comparing the findings from different protocols.

*Lateralization and localization.* Analysts reported their institutions requested fMRI to lateralize language cortex (100%), and in most cases also to guide surgical margins to avoid language cortex (59%) [n=54]. The location of activation is typically evaluated by overlying the data on the patient’s T1 image (89%), and less frequently on the patient’s T2 (29%) or a canonical T1 (e.g., MNI152; 7%). Four analysts (9%) independently noted overlaying the data on FLAIR imaging for specific lesion types (e.g., dysplasia), and another noted overlaying the data on an average EPI during interpretation to ensure the effects of EPI distortion did not impact interpretation.

*Reporting.* Almost all respondents reported the referrer or team review a written report (82%) and review the images visually at surgical conference (78%) [n=60]. The individual involved in analysis typically interprets this data at the conference (67%). Numerous programs also use the fMRI data in an intraoperative system (e.g., STEALTH) (55%). Laterality indices (LIs) are used at 35% of programs, and an equal proportion of respondents use LIs based on select regions (22%) and whole brain activity (22%).

### 3.5 Practical considerations

*Personnel.* Respondents reported that no single profession is typically responsible for clinical language fMRI, with professionals from at least five queried disciplines completing aspects of fMRI at different sites (TABLE II). At any given program, clinical language fMRI is typically completed by individuals from over two backgrounds (average 2.54, SD 0.9; range 1–5) [n=50 respondents]. When a single individual is responsible for fMRI, they are trained as a neuropsychologist (40%), radiologist (40%), or received doctoral training in neuroimaging (single instance). Most programs completing fMRI involve input from individuals trained as radiologists (66%) and MR technologists (64%), with other professionals frequently involved including neuropsychologists (36%), neurologists (24%), neuroscientists with expertise in physics or engineering (22%), neuroscientists more broadly (12%), general research assistants (20%) and individuals from other professions (10%; e.g., neurosurgery).

**Table II:**
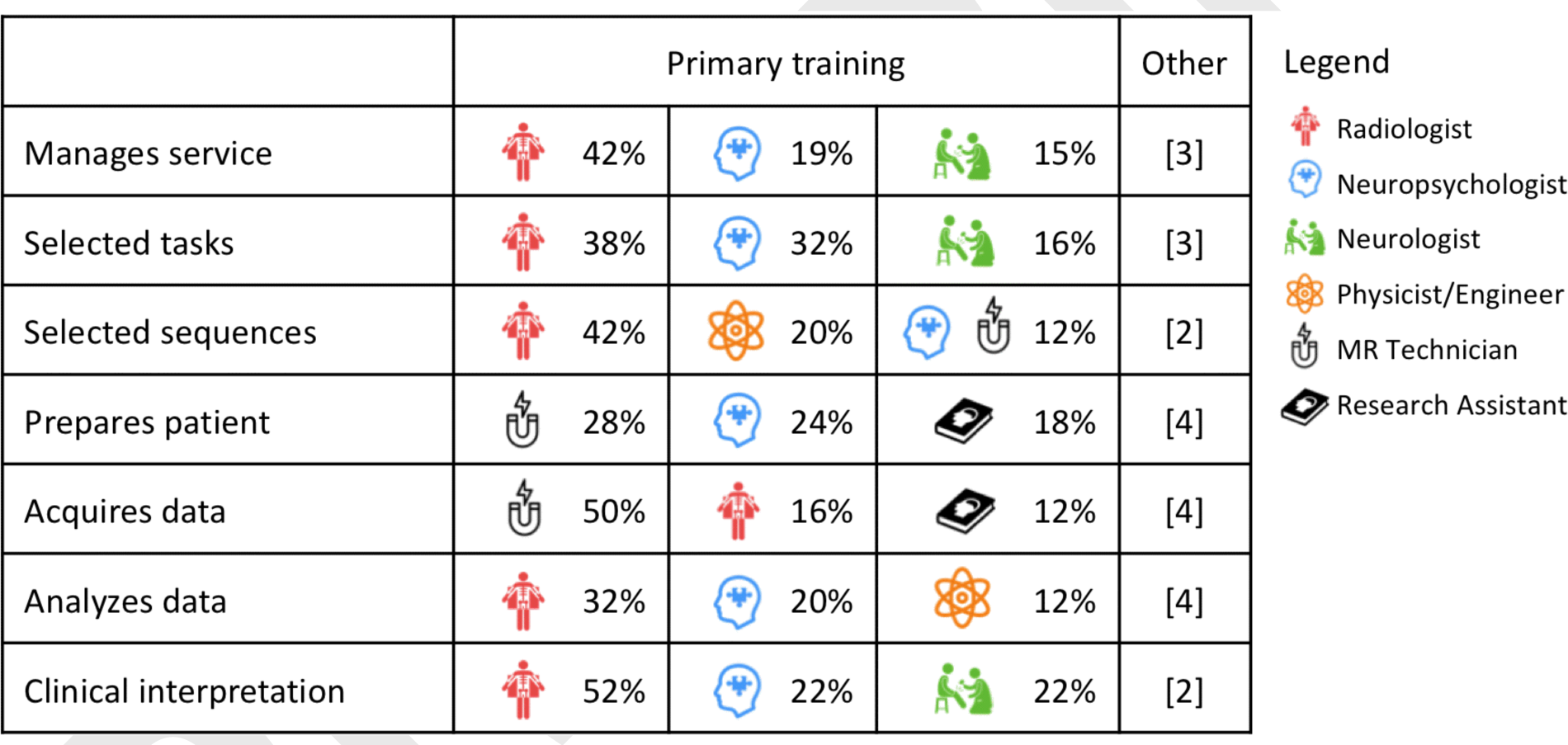
Primary training of individuals responsible for different aspects of clinical language fMRI in presurgical epilepsy programs. (3 most frequently reported disciplines). Other professions are less frequently involved in (i) management [MR technicians; phys./eng.; neuroscientists]; (ii) /task selection [neuroscientists; phys./eng.; MR technicians]; (iii) sequence selection [neuroscientists; neurologists]; (iv) patient preparation [radiologists; phys./eng.; neuroscientists; neurologists]; (v) data acquisition [neuropsychologists; phys./eng.; neuroscientists; neurologists]; (vi) data analysis [research assistants; MR technicians; neurologists; neuroscientists]; and (vii) clinical interpretation [phys./eng.; neuroscientists]. Note that “Physicist/engineer” includes individuals who have this area as their primary training but work as neuroscientists. Additional instances classified as “other” by respondents included a “technician trained in MRI” (sequence selection; patient preparation; analysis); varying professionals (patient preparation); a PhD Engineer who not classified as a neuroscientist (analysis); a neurosurgical research associate (all stages except interpretation); and a neurosurgeon (management of the clinical fMRI service).

*Time and billing.* Analysts reported spending on average 2.3 hours analyzing language fMRI data, including all required tasks (e.g. preprocessing, analysis, thresholding, report generation, exporting to BrainLab etc.) (0.1–11 hours; SD 2.1) [n=47]. Sites who reported hours billed, bill an average of 2 hours (1–8; SD 1.8) for clinical fMRI [n=15]. In the US, details of CPT code use was provided by a fifth of respondents [n=13]. While these response rates limit generalizability, respondents most often billed fMRI as entirely physician or psychologist administered, either with a report (54%; code 96020) or without (46%; code 70555). A third (31%) also bill fMRI time as non-physician/non-psychologist administered (code 70554). Clinical fMRI is also billed as neuropsychological assessment (96118, paired with 70555), and another site billed a diagnostic radiology code (70553; this site also noted billing 70555 and 96020). Three European respondents reported billing fMRI at 300, 1000, and 3500 euro.

*Funding sources.* Language fMRI used to determine surgical eligibility, plan resection or counsel on surgical risk is primarily funded through clinical means (insurer/patient; 86%) [n=44]. Programs who bill fMRI clinically do so for a majority of their patients – 92% (range 20–100; SD 17.4). Approximately a quarter of the responding programs (23%) also or alternately use research funding to pay for fMRI used clinically in 45% of their patients (range 10–100; SD 32.3). Other sources also fund at least some clinical fMRI imaging in a fifth of programs (21%); those funding sources support imaging in 57% of patients at these sites (4–100; SD 46.8). One Canadian site noted no billing code was used.

### 3.6 fMRI accuracy and outcomes

As noted, most analysts reported limited involvement in clinical care, with 65% not involved in surgical decision making at all (65%). While this limits a detailed and unbiased discussion of fMRI relative to clinical outcomes (and clinicians’ reports of outcome are available elsewhere) [Benjamin et al., 2018], note that analysts considered language fMRI to have successfully identified the dominant hemisphere 84% of the time (SD 15%; 20–100%) [n=52 respondents]. fMRI was most often reported to never have disagreed with other methods (54%), or alternately to have disagreed in at least one instance with Wada (32%), stimulation mapping (12%), or other methods (16%) [n=50]. Respondents who could comment often did not know whether, in these cases, fMRI or the other method was correct (42%) [n=26]. Of those who could comment, an equal number reported instances where fMRI was (35%) and was not (35%) judged correct. Of those centers reporting cases of discordance, 17% (4 of 24) had published this finding.

When asked about cases of persistent (>3 month) postoperative language decline when all fMRI-positive language sites were preserved, 44% of respondents did not know about outcomes [n=52]. With the above caveats, of the 29 who did, six (21%) reported instances of decline (two of these cases were also included in the clinician survey) [Benjamin et al., 2018]. None of these cases had been published. A question about whether patients had maintained pre-operative language ability despite resected fMRI-positive language cortex was answered by a subset of respondents who both knew their program did use fMRI to guide surgical margins and avoid language cortex, and knew their program would resect fMRI-positive cortex in some instances [n=14]. Of these, nine (64%) reported cases of maintained function (three of these cases were also included in the previous clinician survey). One (11%) had published this finding.

## 4.0 Discussion

Beyond the specific findings on aspects of task design, acquisition, analysis, and practical issues in clinical language fMRI, the central finding of this work is that virtually all aspects of practice currently vary markedly across programs. Each of the best-validated approaches [Szaflarski et al., 2017] is used by fewer than 10% of programs. As with clinicians, analysts report that their maps are frequently used to guide surgical margins to preserve language function (59% of sites). An important related finding is that clinical programs typically vary statistical thresholds on a patient-by-patient basis (79% of programs). This approach is often seen as both clinically responsible and indeed essential in ensuring no language cortex is inappropriately resected, is emphasized in some clinical guidelines [American College of Radiology, 2014], and may allow more accurate results when the degree of variation in other aspects of fMRI is noted. Conversely, it is also frequently seen as opposing standardization, objectivity, and the research method. Note, however, that the best validation studies to date do actually use a highly standardized approach [Bonelli et al., 2012; Janecek et al., 2013; Szaflarski et al., 2017]. As in a more comprehensive survey of clinicians’ experience of clinical language fMRI [Benjamin et al., 2018], analysts frequently (46%) report that language laterality judged by fMRI has disagreed with that yielded by other methods in at least some instances. Finally, consistent with professional guidelines recommending clinical fMRI involve the multiple professions it requires [Bobholz et al., 2004], these results show the method is typically interdisciplinary. A caveat is that fMRI is frequently completed without input from experts in the assessment of cognition or psychometric design (36% of programs).

These points highlight the descriptive rather than prescriptive nature of the survey results: these findings describe current, and not best, practices in clinical language fMRI. It is likely that in at least some instances the procedures in widespread use reflect reasonable default or historical settings. For example, the most commonly used smoothing kernel in sites using the software package SPM was 8mm (60%) [n=10]; notable as both a reasonable choice and as SPM’s default kernel value. The data here offer a useful starting point in clinical fMRI protocol design, particularly for details not document in published literature, but published evidence and validation studies [Szaflarski et al., 2017] will be of greater use. These findings also highlight the importance of studies of clinical fMRI making clear all detail required for replication (e.g., patient factors; task design, acquisition and analysis; see Supplement 3).

### 4.1 Cognitive design

The observed variation in tasks’ cognitive design will certainly result in different patterns of activation in clinical fMRI across centers, while overall laterality is more likely to remain constant. The range of tasks used likely reflects fMRI’s evolution from an in-house research tool to a clinical method, and insufficient knowledge of how the cognitive task used will influence results. It is also notable that the structure of the three most commonly-used tasks does not center on the cognitive deficit most frequently observed after dominant temporal surgery: naming decline [Sherman et al., 2011]. Further, while fMRI based on verbal fluency (“letter-cued word generation”) has been shown to be predictive of language impairment in some samples [Bonelli et al., 2012], it is notable that dominant temporal resection does not differentially impair this cognitive function and may actually result in its improvement [Sherman et al., 2011]. Large-scale, head-to-head comparisons of protocols are required to develop optimal, standardized, and reproducible tasks and analytic approaches. Maximally-sensitive tasks will likely also use control stimuli that are cognitively matched, as language tasks have been shown to reveal brain regions to differing extents when an active control is used instead of simple “rest” [Binder, 2011]. These data also highlight the need for validation of clinical fMRI in languages other than English. While few such validated protocols were identified in a recent extensive review [Szaflarski et al., 2017], our data show a large number are in use. Simple, validated clinical screening tools to document language proficiency for fMRI are also required.

As in all cognitive assessment, rigidly standardized patient instructions, administration, and evaluation of compliance will be essential to control the cognitive strategy used and brain regions engaged. To allow standardization, this needs to be done with the most challenging patients in mind. Variation even in the wording of instructions (e.g., “relax and be still” vs “relax and ignore the noise”) alters patterns of activation [Benjamin et al., 2010]. The extent of task exposure before scanning (in training) also requires consideration: activation can be reduced when stimuli are repeated [Grill-Spector et al., 2006]. It is also important that the operator directly evaluates the patient’s comprehension. If a task is too complex, or patients are not instructed before scanning, they may experience high anxiety during task blocks or move and alter signal. Confirming the patient completed the task (compliance), as is currently done in 2/3 of the paradigms in use, is also important.

### 4.2 Acquisition and analysis

An area of relative consistency was in acquisition, where nearly all sites acquire images at 3T with isotropic voxels, though the reported variation in acquisition resolution (1.5–4mm) will of course lead to variation in the extent of activation. While preprocessing is largely standardized, many sites use slice-timing (57%) though its use in block designs (used by 95% of respondents) may result in over-smoothing and removal (or introduction) of activation. The optimal degree of smoothing (which varied here from 0–10mm) is also open to debate. One recommendation has been to smooth by the size of the expected activation, though this is less useful when structures of differing sizes may be engaged; another is to smooth by a proportion (e.g., 200%) of the voxel size. Signal drift across time during scanning is a complex issue which some software (e.g., SPM) explicitly models. Removing individual volumes prior to analysis (34% of sites) will distort this modeling and introduce artifact.

Individuals completing clinical fMRI face a choice between open-source, freely available software and closed-source and for-profit commercial alternatives. Closed, prepackaged software typically does not undergo independent review and often lacks transparency in the choice of critical variables and analytic decisions. This gives the incorrect impression that knowledge of image processing, statistical analysis, and cognitive design is not required for fMRI. Further, the impact of these decisions on language maps (and clinical care) can remain opaque. For instance, when asked “how is task-related activation identified?” (general linear modeling/correlation coefficient/other), the three survey respondents who could not answer the question (one noting ‘Bonferroni’) all reported using commercial analysis software. Excellent alternatives that were used in the work validating language fMRI, and which have undergone (and continue to undergo) auditing, debugging, and refinement with recent advances, are readily available in free packages such as SPM, FSL and AFNI.

The neurocognitive model held by the analyst determines the brain regions they expect to see, and thus their results and clinical interpretation. Historic models emphasizing the role of Broca’s and Wernicke’s areas in “expressive” and “receptive” speech exclusively are a useful heuristic, but do not reflect all language-critical areas [e.g., Benjamin et al., 2017; Tremblay and Dick, 2016]. Expressive and/or receptive deficits can follow either anterior or posterior lesions in at least some instances; this is also true to varying degrees in at least four further regions and almost certainly more [Hamberger et al., 2001]. Respondents here reported routinely mapping Broca’s area and Wernicke’s areas (70%), though rarely basal temporal (23%) or other language regions (32%); a cause for consideration if fMRI is used for localization. While such use is not evidence-based, the apparent specificity and precision of fMRI is seductive and lends itself to misinterpretation by even those very familiar with fMRI. In short, a detailed understanding of brain-cognition relationships; the brain regions activated by a given task in a given patient; and the limitations of fMRI are essential in clinical fMRI. These skills allow those conveying fMRI findings to do so confidently with an eye to the utility of their method and its limitations, and guide surgical teams to accurate interpretation of findings.

### 4.3 Clinical language fMRI is equal parts imaging assessment and neurocognitive assessment

Many centers are apparently performing clinical fMRI without all required expertise. Early guidelines set out these skills [Bobholz et al., 2004], noting the importance of (among other skills) knowledge of neuroanatomy; the structure of cognition; MR physics and image artifacts; statistical analysis; and the use and development of psychological tests. Recommended parameters and protocols were, however, absent. More recently guidelines generated through radiology are superior in detailing aspects of how data is collected and analysis is completed. They are silent, however, on the equally essential requirement for expertise in cognitive design and structure-function relationships [American College of Radiology, 2014], which counters the contribution of specific recommendations for imaging parameters.

No discipline currently receives training in all skills required for clinical language fMRI [Bobholz et al., 2004]. Radiologists do not typically have the skills required after standard residency and fellowship training. While expert in MR imaging, they will usually require additional training in cognition, psychometric task design, and functional neurology. Clinical neuropsychologists do not typically have the skills required for clinical language fMRI after typical training. They have expert knowledge in cognitive assessment and brain-behavior relationships, but do not receive training in statistics, MR physics, and clinical imaging essential for fMRI. Neurologists usually have some degree of training in each relevant domain, but lack the required depth of knowledge. Doctoral researchers in fMRI are experienced to varying degrees in cognitive task design, data acquisition, and analysis, but frequently lack knowledge of brain pathology, clinical care in epilepsy, and the integration of this information in surgical decision-making.

While a professional trained in any of the above disciplines may have obtained all the skills required for safe and successful use and interpretation of fMRI, this cannot be assumed. Indeed, as fMRI is fundamentally (i) an imaging assessment and (ii) a neurocognitive assessment used to guide neurosurgery, determine if it is safe to resect brain regions, it is perilous that its use by individuals not credentialed for both cognitive assessment and fMRI occurs. The current professional fragmentation of fMRI likely reflects, to some degree, a need for hospitals to provide fMRI in the absence of a definition of the skills fMRI requires.

### 4.4 What is the correct clinical fMRI protocol?

A model for gold-standard clinical language fMRI already exists in the technology fMRI is largely replacing: the Wada test. Here the requirement for an assessment integrating medical and cognitive assessment led to a formal team-based approach integrating members with the required skills. As the field matures, it is likely that approaches short of this-e.g., run by one individual without a degree of involvement by a complementary profession-will be understood as sub-standard. This structure is already supported to an extent in the US, where multiple professions, including medical doctors and (neuro)psychologists, can and do bill for fMRI.

The ‘correct’ protocol per-se is one based on a published, peer-reviewed study showing that it reliable and valid for the intended purposes (e.g., lateralizing or localizing language; predicting post-surgical decline), in an equivalent patient population. It should be both executed as outlined in that study, i.e. with patients prepared and instructed in a similar manner; equivalent imaging parameters, cognitive design; and analysis; and the results should also be interpreted consistent with the initial study. The method would ideally have also been validated by independent researcher groups in independent samples. If data are not available in a given patient population, the use of a task validated in a similar population may be appropriate until further evidence has accrued. An overview of the evidence for published tasks is available in the American Academy of Neurology guidelines [Szaflarski et al., 2017].

It is important to note that when judging if any given clinical language fMRI has been successful, whether or not a map ‘looks good’ is immaterial. Areas that are language-critical may be entirely absent from a map due to the task’s cognitive design, or how a given patient performed the task. The boundaries of language areas will change with numerous variables. Signal may be missing (e.g., in basal temporal areas) and be misinterpreted as “functionally silent” cortex by a surgical team. The individual analyzing data may be unaware of what brain regions should be shown, how the analysis steps have altered the results, or how the patient’s level of cognitive function has altered the maps.

### 4.5 Limitations

A key limitation of this work is our inability to accurately link given protocols to cognitive outcomes. While we attempted this, the length and complexity of the surveys likely hindered coordination here. Regardless, these data provide insight into current consistencies and inconsistencies in clinical fMRI. We also failed to sample all aspects of fMRI that may influence results; for instance, the number of variables of a task modeled can decrease explained variance. Our use of Snowball Sampling – contacting prominent organizations and individuals, then asking that they identify and invite others to take part and forward the survey to their colleagues (and so on) – helps increase the number of included responses, but precludes calculation of an exact response rate estimate. Within the US we directly contacted 221 NAEC programs, suggesting a low response rate of approximately 13%. In spite of this, the overall sample (63) compares well with other recent surveys [e.g., n=56; Hamberger et al., 2014]. Conversely, these data disproportionately reflect US programs (44%), and multiple large regions (notably in Asia) are under-represented. Future surveys may obtain a broader geographic sample through acquiring data at major conferences, where representatives from programs world-wide can be easily engaged. These data also reflect practice in high volume, academic programs where a majority (65%) of patients receive language fMRI. We might expect the heterogeneity observed would only increase with greater representation of smaller and non-academic surgical programs.

### 4.6 Summary

These findings constitute the first comprehensive description of language fMRI in the clinic. They suggest a marked split between the evidence supporting fMRI’s use and its clinical implementation, and that standardization of the optimal protocols, analysis, and skills required for required for successful fMRI is much needed. Taking these steps will allow the field to converge on a standardized and optimal approach to provide the best patient care.

## Acknowledgements

Thank you to our respondents, who gave the significant time and energy required for this survey.

This work was supported by Yale CTSA [UL1TR000142] from the National Center for Advancing Translational Science (NCATS), National Institutes of Health USA; and the Swebilius Foundation.

Richard A. Bronen discloses he is a consultant for Pfizer, Inc. None of the authors reports a conflict of interest.

## SUPPLEMENTARY MATERIAL

Supplement 1: Survey.

Supplement 2: Full responses regarding cognitive task design, as summarized in Table I.

Supplement 3: Note on FDA approval and fMRI software.

